# Artificial light at night and an invasive snail synergistically enhance the invasion of a non-native macrophyte

**DOI:** 10.64898/2025.12.01.691521

**Authors:** Jingjing Xue, Ayub M.O. Oduor, Hong-Li Li, Feng Li, Yanjie Liu

## Abstract

Herbivory shapes plant invasion outcomes, yet its role in aquatic plant invasions under changing environmental conditions, such as artificial light at night (ALAN), remains poorly understood. We conducted three experiments using invasive macrophyte *Myriophyllum aquaticum*, a native macrophyte community (*Vallisneria natans*, *Hydrilla verticillata*, *M. spicatum*), invasive snail *Pomacea canaliculata*, and native snail *Cipangopaludina chinensis* to test the combined effects of ALAN and herbivory on non-native macrophyte invasions. ALAN increased *M. aquaticum* height and total biomass, but had no effect on native species growth. In feeding assays, *P. canaliculata* consumed all three native species but consistently avoided *M. aquaticum* under both light treatments. *C. chinensis* showed no feeding in no-choice assays, but in choice assays, consumed *H. verticillata* and *M. spicatum* under No-ALAN and only *M. spicatum* under ALAN. In community mesocosms, *P. canaliculata* reduced native macrophyte biomass by 48.0% under No-ALAN and 87.2% under ALAN without affecting *M. aquaticum*. This selective feeding increased *M. aquaticum*’s proportional biomass, with a stronger effect under ALAN than No-ALAN. These results suggest that ALAN can indirectly facilitate non-native macrophyte invasion by amplifying their relative biomass within native communities, particularly in the presence of invasive herbivores, and may promote invasional meltdown through altered feeding preferences.

## 1 | INTRODUCTION

Understanding the factors that govern invasion success of non-native plants is critical for predicting and managing invasions. Herbivory, a key biotic interaction, can strongly influence invasion outcomes. According to the enemy release hypothesis, non-native plants gain a competitive advantage in introduced ranges by escaping their co-evolved specialist herbivores (Keane & Crawley 2002). However, even without these specialists, non-native plants frequently encounter herbivory from native or non-native generalists (Parker et al. 2006). The outcomes of these herbivore-plant interactions are highly variable: some studies show that well-defended non-native plants resist herbivores more effectively than native plants, while others reveal greater susceptibility to herbivory (Meijer et al. 2016). In some cases, non-native invasive plants support a higher diversity and biomass of herbivores and experience greater herbivory, yet continue to dominate communities, likely because their rapid growth rates offset the damage (Allen et al. 2021). Occasionally, invasive plants have been introduced together with their co-evolved herbivores, fostering facilitative interactions that accelerate invasion in line with the invasional meltdown hypothesis (Simberloff & Holle 1999; Parker et al. 2006). For example, non-native snails can facilitate the invasion of non-native macrophytes by preferentially feeding on co-occurring native macrophytes, thereby weakening biotic resistance and giving the invaders a competitive advantage (Yan et al. 2024). Although studies show that herbivory can either suppress or facilitate plant invasions depending on herbivore preferences and plant traits, most research has focused on terrestrial systems (Meijer et al., 2016). Despite freshwater herbivores consuming 40-48% of plant biomass compared to just 4-8% in terrestrial ecosystems (Bakker et al. 2016), we know little about how herbivory affects aquatic plant invasions under changing environmental conditions (Yan et al. 2024).

Artificial light at night (ALAN) is a growing anthropogenic disturbance that alters physiology, growth, and species interactions in both terrestrial and aquatic organisms (Hölker et al. 2023). In terrestrial plants, ALAN can reduce net photosynthetic rates in species (Wei et al. 2023), and modify biomass and traits linked to growth and stress resistance (Bucher et al., 2023). Plant responses to ALAN can vary with their invasive status. For example, under ALAN, invasive plants gained more biomass than natives (Murphy et al., 2021; Liu et al., 2022; Kawawa Abonyo & Oduor, 2024). Urban and suburban water bodies typically experience ALAN of 0.1-20 lux (Jechow & Hölker 2019), with some highly illuminated areas reaching up to 150 lux (Bolton et al. 2017). In freshwater plants, ALAN alters leaf physiology and chemistry, affecting photoprotection, photosynthesis, and flowering, with cascading effects on competition and herbivory (Segrestin et al. 2021). ALAN also influences traits such as leaf toughness, nutrient content, and defensive compounds (e.g., tannins, phenolics, terpenes) that determine palatability to herbivores (Cao et al. 2024). For invertebrate herbivores, ALAN shifts diel activity, foraging efficiency, and habitat use (Ganguly & Candolin 2023), with cascading effects on plant-herbivore interactions (Cieraad et al. 2022). Yet, how ALAN and herbivory interact to influence aquatic plant invasions remains poorly understood (Liu & Heinen, 2024).

Herbivory intensity is shaped by plant palatability, which depends on structural traits (e.g., tissue toughness, leaf density) and chemical defenses (e.g., tannins, phenolics, lignin) (Carmona et al. 2010). Light availability regulates these traits by controlling the biosynthesis of structural and secondary compounds; low light can reduce lignin and toughness, increasing susceptibility to herbivores, whereas high irradiance can enhance defenses (Chen et al. 2002; Cao et al. 2024). Aquatic habitats are naturally dim, with light levels reduced to 1–10% of surface values within the first meter (Hölker et al. 2023), and submerged macrophytes are adapted to these conditions, achieving photosynthetic saturation at just 0.5-3% of surface light (Chen et al. 2020). ALAN may disrupt these adaptations, altering plant palatability and herbivore feeding patterns, potentially driving invasional meltdown if invasive herbivores preferentially target natives. However, the combined effects of ALAN and herbivory on the invasion of non-native macrophytes into native communities have not been tested.

To address this knowledge gap, we conducted three complementary experiments using the invasive macrophyte *Myriophyllum aquaticum*, a native macrophyte community (*Vallisneria natans*, *Hydrilla verticillata*, *Myriophyllum spicatum*), the invasive snail *Pomacea canaliculata*, and the native snail *Cipangopaludina chinensis*. Our research was structured around three sequential hypotheses. First, to establish a baseline effect in the absence of herbivory, we hypothesized that ALAN promotes greater biomass accumulation in *M. aquaticum* than in native macrophyte species, consistent with findings in terrestrial systems. Second, we tested the hypothesis that ALAN reduces macrophyte palatability, thereby lowering consumption by both native and invasive snails and decreasing grazing pressure on the invasive species. Finally, we tested whether ALAN and herbivory interact to amplify the competitive dominance of *M. aquaticum* over native macrophytes. Collectively, these experiments evaluate the overarching hypothesis that light pollution can trigger invasional meltdown, whereby multiple invaders act synergistically to accelerate native community decline.

## 2 | MATERIALS AND METHODS

### 2.1 | Study system

We conducted three mesocosm experiments at the Dongting Lake Wetland Station, Hunan, China (29°30′N, 112°48′E), within a subtropical monsoon climate (mean annual temperature 16.8 °C; precipitation 1,305 mm) characteristic of the Yangtze River floodplain (Xiong et al. 2025). The region, increasingly affected by light pollution from urbanization (Xuan et al. 2024), supports diverse macrophyte communities. Our focal plants included the non-native invasive *M. aquaticum* (introduced from South America in the 1980s; Xiong et al. 2021) and three co-occurring natives: *V. natans*, *H. verticillata*, and *M. spicatum* (**Appendix S1**: **Table S1**). Although treated as natives in this study, *H. verticillata* and *M. spicatum* are among the world’s most problematic aquatic invaders, with major ecological impacts reported in North America, Europe, and Australasia (Moody and Les 2002; Gu 2006). This dual perspective allows us to interpret the growth responses of *H. verticillata* and *M. spicatum* under ALAN and herbivory not only within the context of local Chinese communities where they are native, but also within a broader invasion biology framework, where the same traits expressed in their native ranges may help explain their invasive success abroad. Herbivore treatments used juveniles of the invasive golden apple snail *P. canaliculata* and the native snail *C. chinensis*, both locally abundant and known to feed on all four species (Yan et al. 2024; **Appendix S1: Section S1**).

### 2.2 | Experiment set ups

#### Experiment 1: Does ALAN promote greater biomass accumulation in M. aquaticum than in native macrophyte species?

We conducted a 50-day monoculture experiment (May 4 – June 24, 2023) without herbivory, using a factorial design with four macrophyte species grown individually under ALAN and No-ALAN (**Appendix S1**: **Figure S1**). We established six zones beside a pond: three illuminated with ALAN from solar lamps (29.33 ± 1.58 lux; 0.36 ± 0.068 μmol mL² sL¹, mean ± SE) and three controls exposed only to ambient light. On May 4, three 5-cm seedlings of a single species were planted into PVC pipes (10 cm diameter × 30 cm height) containing a 3-cm layer of 1:1 sieved topsoil and river sand, with 28 cm water depth. All pipes were covered with 1.5-mm mesh to exclude herbivores. We prepared 20 pipes per species (80 in total) and randomly assigned them to the six zones, yielding 10 replicates per treatment. At harvest, plant height and dry biomass (oven-dried at 65 °C) were measured. Due to mortality, 77 pipes were analyzed.

#### Experiment 2: Does ALAN reduce macrophyte palatability and herbivory on M. aquaticum?

We conducted choice and no-choice feeding bioassays with four macrophyte species grown under ALAN and No-ALAN conditions (**Appendix S1**: **Figure S2**). Similar-sized *P. canaliculata* and *C. chinensis* (shell height 1.5–2.5 cm) were starved for 24 h prior to trials to standardize hunger levels. In no-choice assays, we placed 10 g of fresh tissue (stems and leaves) from a single macrophyte species into individual 2-L plastic buckets containing 11 cm of groundwater. In choice assays, 10 g of each of the four macrophyte species (totaling 40 g) was combined per bucket. All plants in a bucket came from the same light treatment. On October 30, one snail was added per bucket, and all buckets were covered with 0.75-mm mesh to prevent escapes and exclude other herbivores. Buckets were then assigned to ALAN or No-ALAN conditions during the bioassay. Treatment combinations (snail species × feeding assay × light) were replicated five times. After four days (November 3), we removed the snails and dried the remaining macrophyte tissue at 65 °C to constant weight. We calculated biomass consumed by first establishing initial dry weights from separate macrophyte individuals not used in the feeding assays. Specifically, we dried 10 g samples (in triplicate) to determine baseline dry masses for each species: *V. natans* (0.453 g ± 0.005 SE), *H. verticillata* (0.562 g ± 0.059 SE), *M. spicatum* (0.430 g ± 0.008 SE), and *M. aquaticum* (1.212 g ± 0.057 SE), following the methodology recommended by Peterson and Renaud (1989). Biomass consumed was then calculated as the difference between the initial and residual dry biomass.

#### Experiment 3: Do ALAN and herbivory interact to enhance M. aquaticum dominance?

We conducted a 116-day outdoor mesocosm experiment in a fully crossed design with light (ALAN vs. No-ALAN) and herbivory (no-snail control, herbivory by *C. chinensis*, or *P. canaliculata*) as factors (6 treatments × 10 replicates = 60 mesocosms; **Appendix S1**: **Figure S3**). Mesocosms were 160-L buckets (56 × 70 cm) containing 15 cm of 1:1 sieved topsoil and sand, filled to 25 cm water depth. On April 8, 2023, we transplanted 16 plants per bucket: four individuals each of *M. aquaticum* (13-cm stems), *V. natans* (seedlings), *H. verticillata* (13-cm stems), and *M. spicatum* (13-cm stems). Dead seedlings were replaced between April 13–15 to ensure full stocking. Light treatments began April 15 with ALAN providing 29.33 ± 1.58 lux at the water surface. All buckets were covered with 0.75-mm mesh to exclude unwanted herbivores. Herbivore treatments were applied in three grazing periods (June 2–8, June 21–July 9, July 28–August 2) by adding six snails (1–2 cm shell height) per bucket, with dead snails replaced to ensure constant herbivore density and sustained grazing pressure as the macrophytes grew. On 2 August, all plants were harvested, sorted by species, and oven-dried (65 °C) to constant weight. Due to macrophyte mortality in three mesocosms, we analyzed dry biomass from 57 buckets. From the dry biomass data, we calculated: (1) total biomass of the native macrophyte community (sum of *V. natans*, *H. verticillata*, and *M. spicatum* biomass); and (2) proportion of *M. aquaticum* biomass relative to total macrophyte biomass per bucket.

### 2.3 | Statistical analysis

All statistical analyses were conducted in R (version 4.3.3; R Core Team, 2024) using dry biomass data. Analytical approaches were tailored to the specific design and objectives of each experiment, as described below.

#### Experiment 1: Macrophyte growth in monoculture without herbivory

We fitted linear mixed-effect models to test whether invasive and native macrophyte species differed in their growth responses to ALAN in the absence of herbivory. Response variables were height (log-transformed), total dry biomass, and root mass fraction (both square-root transformed). Root mass fraction (root/total biomass) indicates biomass allocation strategy, with higher values reflecting greater belowground investment and lower values reflecting greater aboveground allocation. Fixed effects were light treatment, macrophyte species, and their interaction, with zone as a random effect. We ensured homoscedasticity by modeling variances per species using the *varIdent* function (Pinheiro & Bates 2023). We assessed the significance of fixed effects with log-likelihood ratio tests (Zuur et al. 2009) and explored significant interactions with Tukey post hoc comparisons using *emmeans* (Lenth 2024).

#### Experiment 2: Herbivore feeding assays

For both no-choice and choice assays, we fitted linear mixed-effect models with biomass consumed as the response variable and light treatment (ALAN vs. No-ALAN), herbivore origin (invasive vs. native snail), macrophyte species, and their interactions as fixed effects. Biomass data were Yeo-Johnson transformed (Kuhn 2008) to meet normality, and variance structures were modeled with *varIdent* function. We included zone and bucket identity (nested within zone) as random factors. The significance of fixed-effect independent variables was assessed using log-likelihood ratio tests.

#### Experiment 3: Interactive effects of ALAN and herbivory on the invasion of M. aquaticum into native macrophyte communities

We used linear models to test effects of light (ALAN vs. No-ALAN) and herbivory (no-snail, invasive snail, native snail) on *M. aquaticum* biomass (log-transformed), native community biomass, and proportional biomass of *M. aquaticum* (both cube-root transformed). To specifically test differences between the effects of native and invasive snails on macrophyte biomass, we created two *a priori* contrasts to break down the herbivore treatment: one comparing the absence of snails to the presence of native snails (Hn), and the other comparing the absence of snails to the presence of invasive snails (Hi). The significance of these effects was assessed using Type II Wald chi-square tests through the *Anova* function in the *car* package (Fox & Weisberg 2019).

## 3 | RESULTS

### Experiment 1: Macrophyte growth in monoculture without herbivory

Macrophyte height and biomass were significantly influenced by species identity and its interaction with light treatment (**Figure 1A-B**; **Appendix S1**: **Table S2**). Post hoc tests showed that ALAN significantly increased the height (36.8%) and total biomass (58.3%) of the invasive *M. aquaticum* compared to No-ALAN, whereas the three native species (*V. natans*, *H. verticillata*, *M. spicatum*) were unaffected (**Figure 1A–B**). The native *V. natans* had a significantly higher root mass fraction than the other macrophytes, and across species, ALAN reduced root mass fraction by 13.0% (**Figure 1C**; **Appendix S1**: **Table S2**).

**Figure 1.**
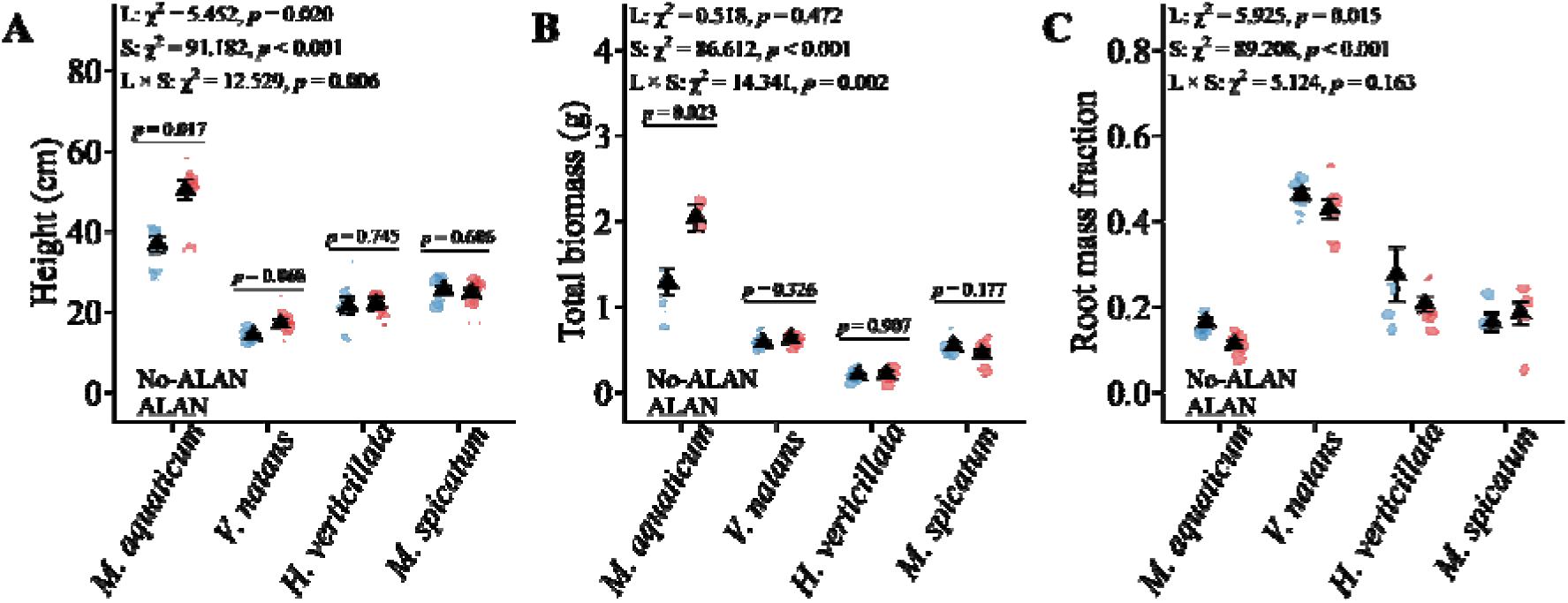
Mean (±1 SE) height (A), total biomass (B), and root mass fraction (C) of the invasive macrophyte *Myriophyllum aquaticum* and three native species (*Vallisneria natans*, *Hydrilla verticillata*, *Myriophyllum spicatum*) grown in monoculture under ALAN and No-ALAN conditions. Black triangles show treatment means; points represent individual replicates. Full model results are provided in Appendix S1: Table S2.

### Experiment 2: Herbivore feeding assays

In both feeding assays, the invasive snail *P. canaliculata* consumed all three native macrophyte species (*V. natans*, *H. verticillata*, *M. spicatum*) but avoided the invader *M. aquaticum*, regardless of light treatment (**Figure 2A & B**; **Appendix S1**: **Table S3**). By contrast, the native snail *C. chinensis* showed context-dependent feeding (significant L × H × S interaction; **Figure 2A & B**; **Appendix S1**: **Table S3**). In no-choice assays, *C. chinensis* did not consume any macrophyte species under either ALAN or No-ALAN condition (**Figure 2A**). However, in choice assays, *C. chinensis* fed on *H. verticillata* and *M. spicatum* under No-ALAN and only on *M. spicatum* under ALAN (**Figure 2B**).

**Figure 2.**
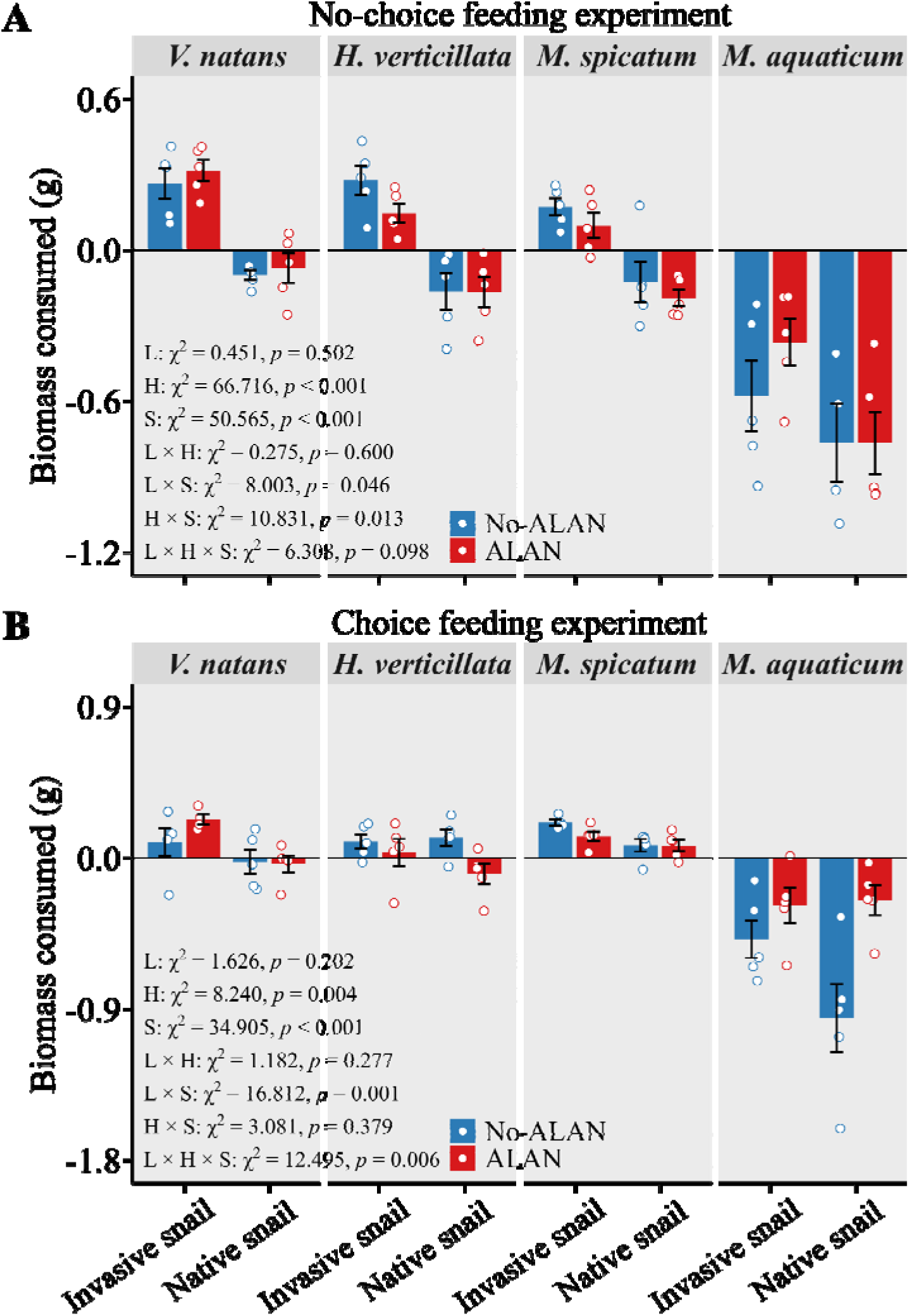
Mean (±1 SE) biomass of native macrophytes (*Vallisneria natans*, *Hydrilla verticillata*, *Myriophyllum spicatum*) and the invasive non-native macrophyte *Myriophyllum aquaticum* consumed by invasive (*Pomacea canaliculata*) and native (*Cipangopaludina chinensis*) snails in no-choice (A) and choice (B) feeding experiments. White points with blue or red outlines indicate individual replicates. Main and interactive effects of light (L), herbivore origin (H), and macrophyte species (S) are shown. Full model results are provided in Appendix S1: Table S3. Since macrophytes can grow in water, residual biomass may exceed initial amounts; thus, zero or positive values indicate herbivore consumption, while negative values indicate none.

### Experiment 3: Interactive effects of ALAN and herbivory on the invasion of M. aquaticum into native macrophyte communities

Relative to the no-herbivory control, neither the native snail *C. chinensis* nor invasive snail *P. canaliculata* significantly affected *M. aquaticum* biomass under either light treatment (**Figure 3A**; **Appendix S1**: **Table S4**). Likewise, herbivory by *C. chinensis* did not alter native community biomass (**Figure 3B**; **Appendix S1**: **Table S4**). In contrast, *P. canaliculata* exerted strong selective herbivory, reducing native community biomass by 48.0% under No-ALAN and 87.2% under ALAN (**Figure 3B**; **Appendix S1**: **Table S4**). This suppression of native macrophytes by *P. canaliculata* consequently increased the proportional biomass of *M. aquaticum* within the community, with effects of 29.8% under No-ALAN and 87.7% under ALAN (**Figure 3C**; **Appendix S1**: **Table S4**).

**Figure 3.**
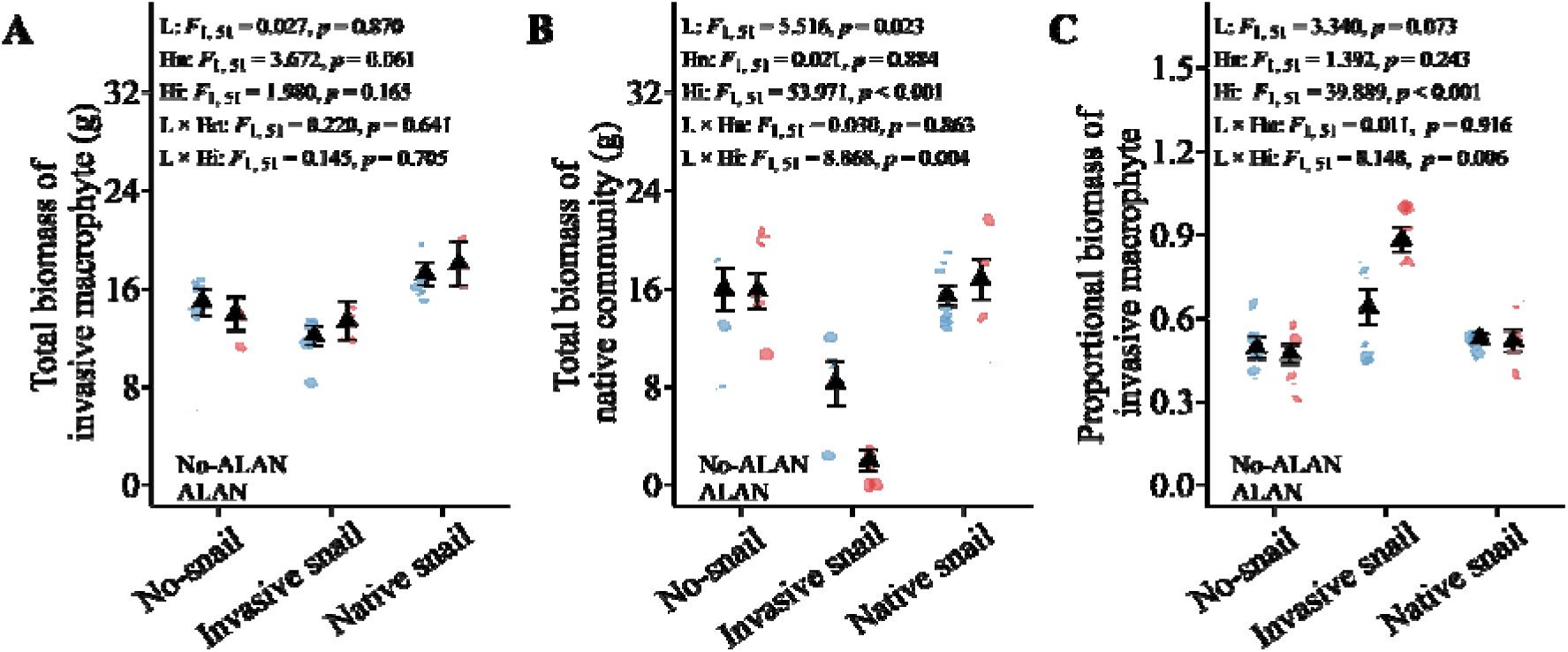
Mean (±1 SE) total biomass of an invasive macrophyte *Myriophyllum aquaticum* (A), a native macrophyte community (*Vallisneria natans*, *Hydrilla verticillata*, and *Myriophyllum spicatum*) (B), and the proportional biomass of *M. aquaticum* within the community (C) under factorial combinations of light (ALAN vs. No-ALAN) and herbivore treatments (no-snail, invasive snail, native snail). Black triangles show treatment means; white points with blue or red outlines indicate individual replicates. Model terms: L = light treatment; Hn = no-snail vs. native snail; Hi = no-snail vs. invasive snail. Full statistics are in Appendix S1: Table S4.

## 4 | DISCUSSION

This study demonstrates that ALAN can drive aquatic plant invasions both directly, by enhancing invasive macrophyte growth, and indirectly, by altering herbivore behavior and amplifying the selective feeding of invasive snails on native communities. A striking paradox emerged: although *H. verticillata* and *M. spicatum* rank among the world’s most problematic aquatic invaders (Moody and Les 2002; Gu 2006), they showed no growth enhancement under ALAN and remained highly vulnerable to herbivory in their native range, losing 48–87% of biomass to *P. canaliculata* and partially to *C. chinensis*. By contrast, *M. aquaticum* combined strong ALAN-driven growth (58% biomass increase) with complete herbivore resistance, reinforcing its dominance through the synergistic effects of light pollution and selective herbivory. These results suggest that the global invasiveness of *H. verticillata* and *M. spicatum* may rely more on release from natural enemies than on inherent competitive superiority, underscoring that ‘invasiveness’ is not an intrinsic trait but an emergent property of ecological context. Overall, ALAN functioned as a powerful catalyst, accelerating *M. aquaticum*’s advantage and restructuring native macrophyte communities.

### ALAN enhances invasive macrophyte growth and alters biomass allocation

Our monoculture experiment revealed species-specific growth responses to ALAN, with the invasive *M. aquaticum* showing significant increases in height (36.8%) and total biomass (58.3%), while native species were unaffected (**Figure 1A & B**). ALAN also altered belowground allocation, reducing root mass fraction by 13.0% across species, although the native *V. natans* maintained a significantly higher root mass fraction than the others (**Figure 1C**). These results suggest that ALAN may confer advantage to invasive macrophytes through both enhanced above-ground growth and altered resource allocation, consistent with terrestrial studies showing greater invader performance under light pollution (Murphy et al. 2022; Kawawa Abonyo & Oduor 2024). It is worth noting that our 30 lux treatment represents the higher end of urban light pollution, making these findings particularly relevant for understanding impacts in heavily urbanized aquatic systems. The enhanced growth of *M. aquaticum* under ALAN, coupled with the general reduction in root investment across species, may be attributed to extended photosynthetic periods and shifted resource allocation from below-ground to above-ground tissues, as observed in other aquatic plants exposed to prolonged photoperiods (Nakagawa-Lagisz & Lagisz 2023; Hussner et al. 2014).

### Contrasting herbivory by invasive and native snails under ALAN

Feeding assays revealed clear behavioral differences between invasive and native snails, with important consequences for macrophyte community dynamics. The invasive *P. canaliculata* consistently consumed all three native species while completely avoiding the invader *M. aquaticum*, regardless of light treatment (**Figure 2A & B**). This pattern of broad consumption but targeted avoidance confirms *P. canaliculata’s* role as a strong generalist herbivore (Qiu & Kwong 2009) and likely reflects strong chemical defenses or low palatability of *M. aquaticum* (Viveros-Legorreta et al. 2022). By contrast, the native *C. chinensis* showed context-dependent herbivory. In no-choice assays, it consumed none of the macrophytes, suggesting low feeding pressure compared to the invasive snail (**Figure 2A**). However, in choice assays, its feeding shifted with light conditions: under No-ALAN it consumed *H. verticillata* and *M. spicatum*, whereas under ALAN it consumed only *M. spicatum* (**Figure 2B**). Similar ALAN-induced shifts in herbivory have been reported for the pond snail *Lymnaea stagnalis*, which increased grazing on *Ceratophyllum demersum* under light pollution (Mondy et al. 2021), likely due to altered activity, foraging efficiency, or plant palatability (Seymoure et al. 2023). The consistent avoidance of *M. aquaticum* by both snails supports the enemy release hypothesis, which argues that invaders gain an advantage by escaping strong herbivory in their introduced range (Keane & Crawley 2002). Overall, the invasive snail’s heavy consumption of native macrophytes and the native snail’s altered preferences under ALAN create a synergistic threat: weakening native plant assemblages while further promoting the dominance of invasive macrophytes.

### ALAN amplifies invasional meltdown between M. aquaticum and P. canaliculata

Our community assembly experiment suggests that ALAN can amplify an invasional meltdown between *M. aquaticum* and *P. canaliculata*. Although neither snail species directly reduced *M. aquaticum* biomass, the invasive snail’s selective herbivory on native macrophytes indirectly facilitated its dominance (**Figure 3A-C**). ALAN intensified this effect nearly three-fold, with *M. aquaticum*’s proportional biomass increasing by 87.7% under ALAN versus 29.8% under No-ALAN (**Figure 3C**). Our supplementary experiments further showed that ALAN accelerated *P. canaliculata* egg hatching and increased hatchling numbers by 28.6% (**Appendix S2**: **Figure S1**), indicating that by boosting population growth, ALAN enables *P. canaliculata* to exert stronger and more sustained herbivory pressure on native macrophytes, which in turn indirectly promotes *M. aquaticum* dominance. This positive feedback loop between the two invaders exemplifies invasional meltdown framework (Simberloff & Von Holle 1999). Comparable reproductive gains under ALAN have been reported in other invasive animals, such as earlier egg-laying and increased reproductive output in *Anolis* lizards (Thawley & Kolbe 2020), likely due to circadian disruption of reproductive physiology (Stanton & Cowart 2024). Together, these results suggest that light-polluted aquatic habitats may be particularly vulnerable to invasion, supporting the "invasion windows" theory that certain environmental conditions can enhance invader establishment (Johnstone 1986).

## Conclusion

Our findings demonstrate that ALAN can act as a powerful driver of aquatic invasions by simultaneously enhancing invasive macrophyte growth and altering trophic interactions. Under ALAN, the invasive non-native snail *P. canaliculata* preferentially consumed native macrophytes while avoiding the co-occurring invader *M. aquaticum*, thereby indirectly reinforcing *M. aquaticum*’s dominance and initiating an invasional meltdown. By boosting the performance of both invasive plants and invasive consumers, ALAN creates a ‘perfect storm’ that accelerates native community decline and reshapes ecosystem structure.

## ACKNOWLEDGEMENTS

Y.L. acknowledges funding from the National Natural Science Foundation of China (W2411021). H.L. acknowledges funding from the National Key Research and Development Program of China (2021YFC2600400). F.L. acknowledges funding from National Natural Science Foundation of China (42171062).

## Appendix S1

### Section S1

#### Study species

##### Macrophytes

The study included the invasive non-native macrophyte *Myriophyllum aquaticum* and three co-occurring natives: *Vallisneria natans*, *Hydrilla verticillata*, and *Myriophyllum spicatum*. These species were selected based on their co-occurrence in Chinese freshwater ecosystems (Huang et al. 2024). We confirmed invasive status of the macrophytes in China from the General Information of Invasive Alien Plants of China (Hao & Ma 2023), the Life Form database (Poppenwimer et al. 2023), and the Flora of China (www.efloras.org). We sourced seedlings of *V. natans* and stem segments of the other macrophytes from an aquatic plant propagation base (Jingzhou, Hubei Province, China, 29°49’N, 113°27′E) and acclimated them in pond water under natural environmental conditions prior to experimentation.

While we classify *V. natans*, *H. verticillata*, and *M. spicatum* as native to China for the purposes of this study, we acknowledge that the distinction between invasive and non-invasive species is context-dependent and geographically variable. Notably, *H. verticillata* is considered one of the world’s most problematic aquatic weeds and has become highly invasive in North America, particularly in Florida and the southeastern United States, where it forms dense monocultures that displace native vegetation (Gu 2006). Similarly, *M. spicatum*, though native to Eurasia including China, is regarded as a major invasive species across North America, where it has spread to 45 U.S. states and three Canadian provinces since its introduction in the 1940s (Moody and Les 2002). *Myriophyllum aquaticum*, native to South America and invasive in China since the 1980s, has become invasive on all continents except Antarctica (Orchard 1981, Hussner 2009, Xiong et al. 2021, Yan et al. 2024).

All four species demonstrate traits typical of successful invaders including rapid growth, efficient vegetative reproduction, and broad environmental tolerance. *Myriophyllum aquaticum* spreads rapidly by vegetative fragmentation (Orchard 1981, Yan et al. 2024), while the three species native to China are capable of both sexual and vegetative reproduction. *Hydrilla verticillata* and *V. natans* occur in both eutrophic and oligotrophic lakes, while *M. spicatum* favors eutrophic habitats (Aiken et al. 1979, Chen et al. 2022, Yan et al. 2024). This broad ecological amplitude and reproductive flexibility are characteristics that facilitate invasion when these species are introduced outside their native ranges (Santamaría 2002).We recognize that the traits we examine in this study – such as biomass production and responses to environmental conditions – may contribute to invasive potential regardless of a species’ status in its native range. Therefore, while our experimental design treats these species according to their status in China, our interpretation considers the broader implications of their demonstrated invasive capacities elsewhere.

##### Herbivores

Herbivores included the invasive apple snail *Pomacea canaliculata*, a generalist macrophyte consumer, and the native snail *Cipangopaludina chinensis*. These snails, similar in appearance (**Appendix S1**: **Figure S4**), have distinct ecological roles: *P. canaliculata* is a generalist herbivore primarily consuming macrophytes (Paz et al. 2019), while *C. chinensis* exhibits both grazing and filter feeding behaviors (Crone et al. 2023). *Pomacea canaliculata* is native to the freshwater wetlands of South America but has invaded various Asian countries, including China, since the 1980s (Carlsson et al. 2004, Qiu & Kwong 2009). Both snails co-occur with the focal macrophytes in Chinese freshwater systems (Yan et al., 2024). We collected adult snails from ponds at the National Field Scientific Observation and Research Station of the Dongting Lake Wetland Ecosystem. Prior to experimentation, they were acclimated in pond water for two weeks and fed daily to minimize stress and maintain natural feeding behavior.

In the herbivory assay, we calculated biomass consumed by first establishing initial dry weights from separate macrophyte individuals not used in the feeding assays. Specifically, we dried 10 g samples (in triplicate) to determine baseline dry masses for each species: *V. natans* (0.453 g ± 0.005 SE), *H.verticillata* (0.562 g ± 0.059 SE), *M. spicatum* (0.430 g ± 0.008 SE), and *M. aquaticum* (1.212 g ± 0.057 SE), following the methodology recommended by Peterson and Renaud (1989). Biomass consumed was then calculated as the difference between the initial and residual dry biomass. Instead of estimating consumption from the difference between initial and final fresh mass, we adopted this approach because the fresh-mass is inherently unreliable due to fluctuating water content, which varies with environmental conditions, time of day, and tissue damage caused by herbivory (Körner 2002, Cebrian & Lartigue 2004).

#### Experimental ALAN facility

To simulate ALAN, we installed LED solar projection lamps (HBT-1911, 3.7 V, IP 55, 180 lm, white 6000 K; HEBOTESOLAR, Shenzhen, China). We implemented the lighting treatments (ALAN vs. No-ALAN) by arranging 60 buckets in a grid (65 cm between rows, 90 cm between columns) within an unshaded outdoor pond at the research station. Each bucket was wrapped in aluminum foil tape to block lateral light transmission and randomly assigned to ALAN or No-ALAN treatments (**Appendix 1**: **Figure S3**). Lamps were mounted on wooden sticks 24 cm above the bucket rim and secured with transparent tape. To achieve ecologically relevant light levels, we covered the lamps with three layers of 300-mesh white nylon netting, which reduced illumination in ALAN-treated buckets to 29.33 ± 1.58 lux (0.36 ± 0.068 μmol ml² sl¹, mean ± SE) at the water surface. This intensity is comparable to typical street lighting (Bennie et al. 2016) and to surface waters near illuminated marine structures (Underwood et al. 2017, Bauer et al. 2022). Control buckets (No-ALAN) experienced only natural moonlight. Any skyglow from nearby urban areas was negligible (< 0.1 lux) due to the rural setting. We measured light intensity using an AS813 illuminometer (SMART SENSOR, Guangdong, China) with a resolution of 1 lux (0 – 200,000 lux). Lamps were programmed to automatically switch on at dusk and off at dawn (from about 7:30 p.m. to 5:00 a.m.), although on 12 overcast days they operated for only *ca.* 4 h.

**Table S1.** Species information for the macrophytes and herbivores used in this study.

| Species | Native/Invasive status in China | Family | Native distributional range | Invasive distributional range | Life cycle/ Reproduction |
| --- | --- | --- | --- | --- | --- |
| *Vallisneria natans*<sup>1</sup> | Native | Hydrocharitaceae | Southeast and Eastern Asia | - | Perennial |
| *Hydrilla verticillata*<sup>1</sup> | Native | Hydrocharitaceae | Africa, Asia, Eastern Europe, Australia | North and South America, England, Germany, New Zealand | Perennial |
| *Myriophyllum spicatum*<sup>1</sup> | Native | Haloragaceae | Europe, Asia, and North Africa | North America | Perennial |
| *Myriophyllum aquaticum*<sup>1</sup> | Invasive | Haloragaceae | South America | North America, South Africa, Japan, England, Western Europe, New Zealand, Australia, and China | Perennial |
| *Pomacea canaliculata*<sup>2</sup> | Invasive | Ampullariidae | South America | United States, South-east Asia | Gonochorism; oviparity |
| *Cipangopaludina chinensis*<sup>2</sup> | Native | Ampullariidae | Eastern Asia | North America | Gonochorism; vivipation |

<sup>1</sup>The native and invasive distributional ranges of the species are based on the information from https://powo.science.kew.org/.
<sup>2</sup>The native and invasive distributional ranges of the species are based on the information from https://en.wikipedia.org/.

**Table S2.** Results of linear mixed-effect models that tested main and interactive effects of light treatment (ALAN vs. No-ALAN) and species (*Vallisneria natans*, *Hydrilla verticillate*, *Myriophyllum spicatum* and *Myriophyllum aquaticum*) on height, total biomass, and root mass fraction of four macrophytes. Significant effects (*p* < 0.05) are shown in bold font.

| Fixed effects | Height |  |  | Total biomass |  |  | Root mass fraction |  |  |
| --- | --- | --- | --- | --- | --- | --- | --- | --- | --- |
| | *df* | $\chi^2$ | *p* | *df* | $\chi^2$ | *p* | *df* | $\chi^2$ | *p* |
| Light (L) | 1 | 5.452 | **0.020** | 1 | 0.518 | 0.472 | 1 | 5.925 | **0.015** |
| Species (S) | 3 | 91.182 | **<0.001** | 3 | 86.612 | **<0.001** | 3 | 89.208 | **<0.001** |
| L × S | 3 | 12.529 | **0.006** | 3 | 14.341 | **0.002** | 3 | 5.124 | 0.163 |
| **Random effects** |  | SD |  |  | SD |  |  | SD |  |
| Zone |  | 0.018 |  |  | 0.006 |  |  | 0.013 |  |
| Residual |  | 0.253 |  |  | 0.096 |  |  | 0.113 |  |
| | Marginal $R^2$ | Conditional $R^2$ | | Marginal $R^2$ | Conditional $R^2$ | | Marginal $R^2$ | Conditional $R^2$ | |
| $R^2$ of the model | 0.693 | 0.695 | | 0.912 | 0.912 | | 0.526 | 0.532 | |

*Note:* For the random effect, zone refers to the site where polyvinyl chloride pipes were placed.

**Table S3.**
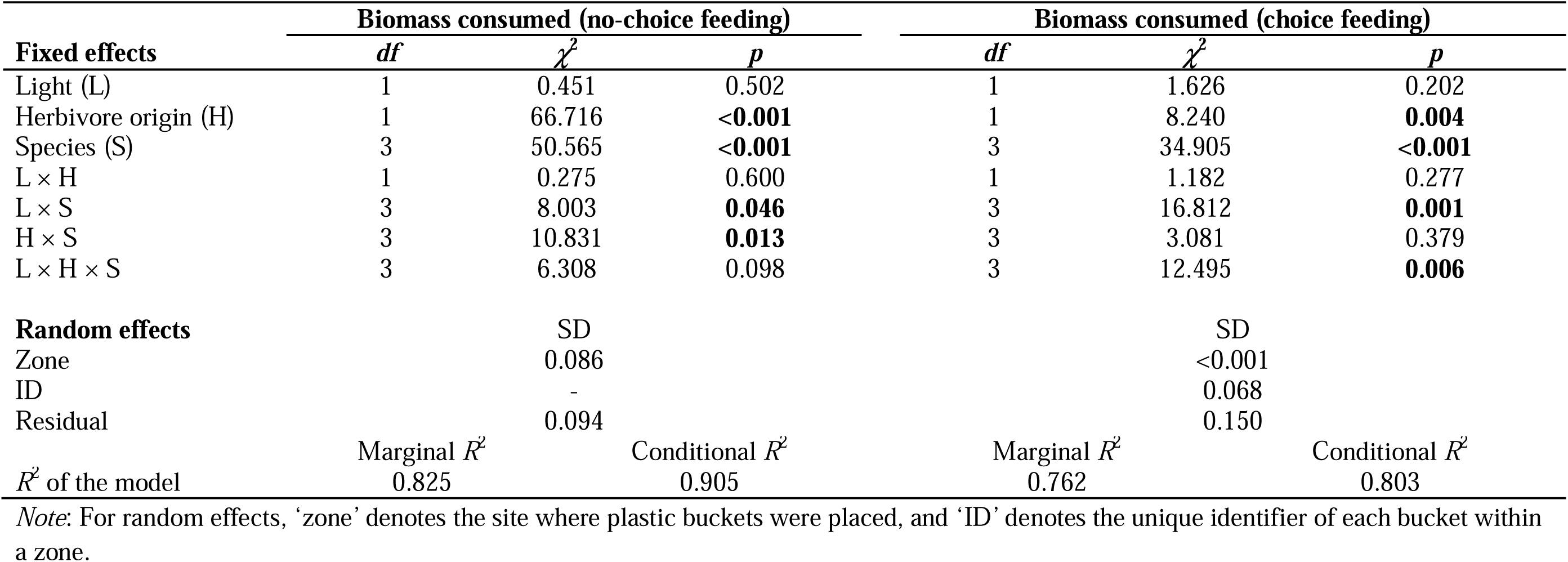
Results of linear mixed-effect models that tested main and interactive effects of light treatment (ALAN vs. No-ALAN), herbivore origin (invasive vs. native snail), and macrophyte species identity (*Vallisneria natans*, *Hydrilla verticillata*, *Myriophyllum spicatum*, *M. aquaticum*) on dry biomass consumed in no-choice and choice feeding experiments. Significant effects (*p* < 0.05) are shown in bold.

| Fixed effects | Biomass consumed (no-choice feeding) |  |  | Biomass consumed (choice feeding) |  |  |
| --- | --- | --- | --- | --- | --- | --- |
| | *df* | $\chi^2$ | *p* | *df* | $\chi^2$ | *p* |
| Light (L) | 1 | 0.451 | 0.502 | 1 | 1.626 | 0.202 |
| Herbivore origin (H) | 1 | 66.716 | **<0.001** | 1 | 8.240 | **0.004** |
| Species (S) | 3 | 50.565 | **<0.001** | 3 | 34.905 | **<0.001** |
| L × H | 1 | 0.275 | 0.600 | 1 | 1.182 | 0.277 |
| L × S | 3 | 8.003 | **0.046** | 3 | 16.812 | **0.001** |
| H × S | 3 | 10.831 | **0.013** | 3 | 3.081 | 0.379 |
| L × H × S | 3 | 6.308 | 0.098 | 3 | 12.495 | **0.006** |
| **Random effects** |  | SD |  |  | SD |  |
| Zone |  | 0.086 |  |  | <0.001 |  |
| ID |  | - |  |  | 0.068 |  |
| Residual |  | 0.094 |  |  | 0.150 |  |
| | Marginal $R^2$ | | Conditional $R^2$ | Marginal $R^2$ | | Conditional $R^2$ |
| $R^2$ of the model | 0.825 | | 0.905 | 0.762 | | 0.803 |

*Note:* For random effects, ‘zone’ denotes the site where plastic buckets were placed, and ‘ID’ denotes the unique identifier of each bucket within a zone.

**Table S4.** Results of linear models and ANOVA testing main and interactive effects of light (ALAN vs. No-ALAN) and herbivore (no-snail, invasive snail, native snail) treatments on total biomass of the invasive macrophyte *Myriophyllum aquaticum*, total biomass of the native macrophyte community, and proportional biomass of *M. aquaticum* within the community. Planned contrasts (Hn: no-snail vs. native snail; Hi: no-snail vs. invasive snail) assessed the specific effects of each snail species. Significant effects (*p* < 0.05) are shown in bold.

| Factor | Total biomass of invasive macrophyte |  |  | Total biomass of native macrophyte community |  |  | Proportional biomass of invasive macrophyte |  |  |
| --- | --- | --- | --- | --- | --- | --- | --- | --- | --- |
|  | *df* | *F*-value | *p* | *df* | *F*-value | *p* | *df* | *F*-value | *p* |
| Light (L) | 1, 51 | 0.027 | 0.870 | 1, 51 | 5.516 | **0.023** | 1, 51 | 3.340 | 0.073 |
| Hn (no-snail vs. native snail) | 1, 51 | 3.672 | 0.061 | 1, 51 | 0.021 | 0.884 | 1, 51 | 1.392 | 0.243 |
| Hi (no-snail vs. invasive snail) | 1, 51 | 1.980 | 0.165 | 1, 51 | 53.971 | **<0.001** | 1, 51 | 39.889 | **<0.001** |
| L × Hn | 1, 51 | 0.220 | 0.641 | 1, 51 | 0.030 | 0.863 | 1, 51 | 0.011 | 0.916 |
| L × Hi | 1, 51 | 0.145 | 0.705 | 1, 51 | 8.868 | **0.004** | 1, 51 | 8.148 | **0.006** |

**Figure S1.**
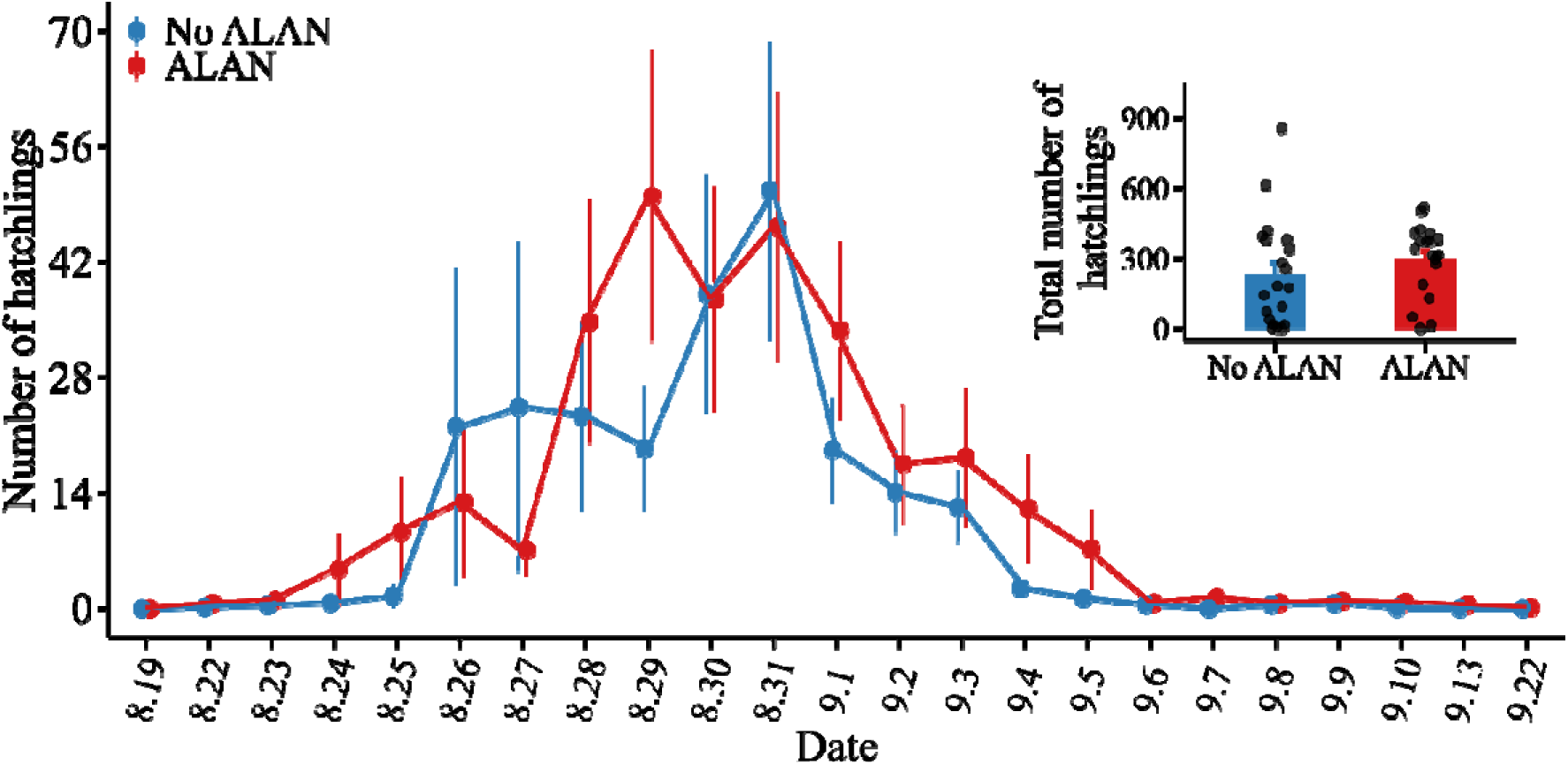
Schematic of Experiment 1 showing polyvinyl chloride pipes with four macrophyte species distributed across ALAN and No-ALAN zones. Forty polyvinyl chloride pipes containing four macrophyte species were randomly distributed across three ALAN zones, with another 40 pipes distributed across three No-ALAN zones (10 replicate pipers per species per light treatment). Each zone contained 13 or 14 randomly placed pipes. Illustrations of macrophytes and light bulbs were obtained from open-source material libraries in WPS Office and combined using Microsoft PowerPoint. The experiment was perfomed at the Dongting Lake Wetland Ecosystem Observation and Research Station.

**Figure S2.**
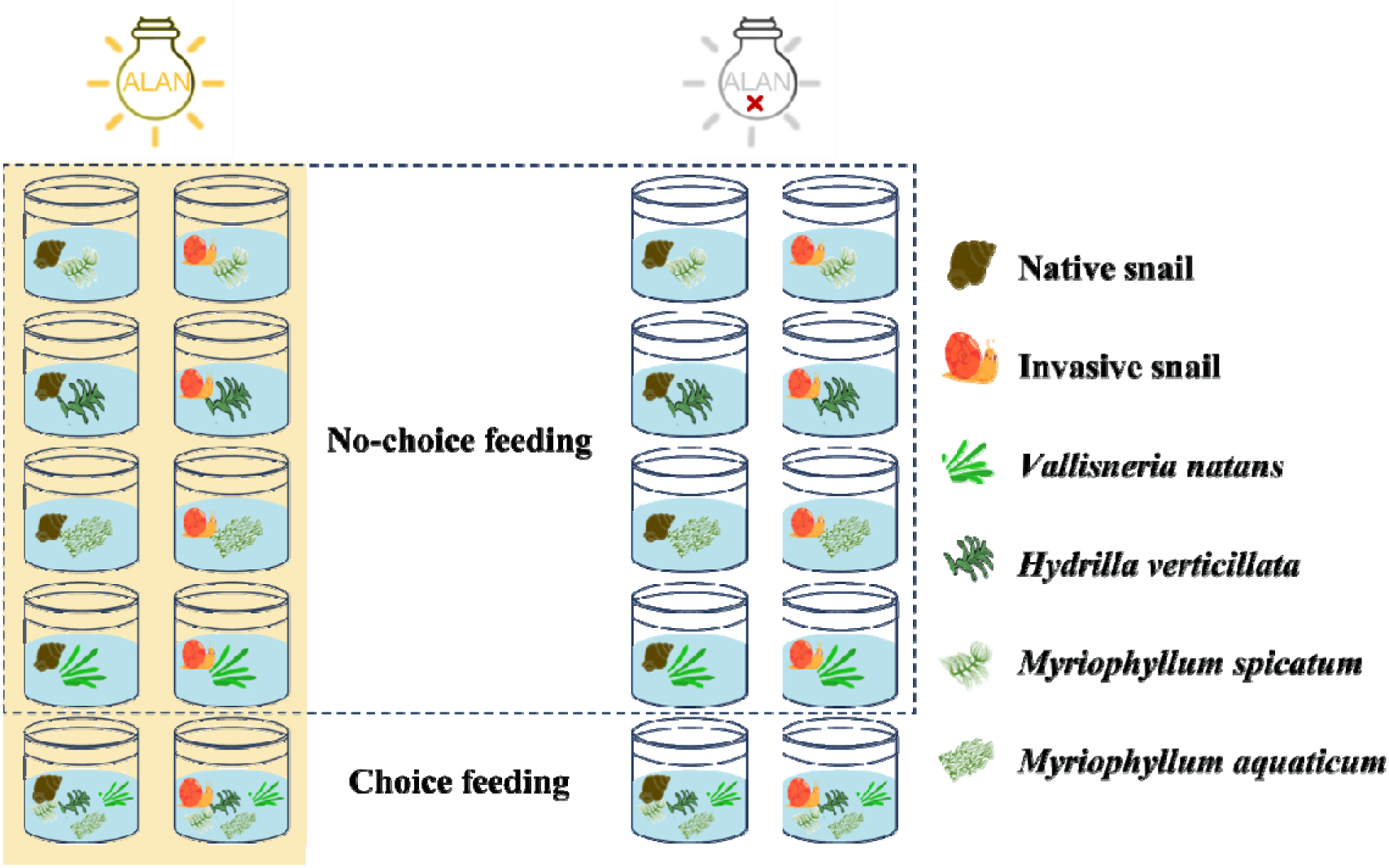
Schematic of Experiment 2 showing no-choice (single species) and choice (four species) feeding assays, crossed with herbivore origin (native *Cipangopaludina chinensis* vs. invasive *Pomacea canaliculata*) and light treatments (ALAN vs. No-ALAN). No-choice feeding assays used tissues from one macrophyte species (*Vallisneria natans*, *Hydrilla verticillata*, *Myriophyllum spicatum*, *M. aquaticum*), while choice feeding assays included all four. Illustrations were adapted from open-source libraries in WPS Office and combined in Microsoft PowerPoint. The experiment was perfomed at the Dongting Lake Wetland Ecosystem Observation and Research Station.

**Figure S3.**
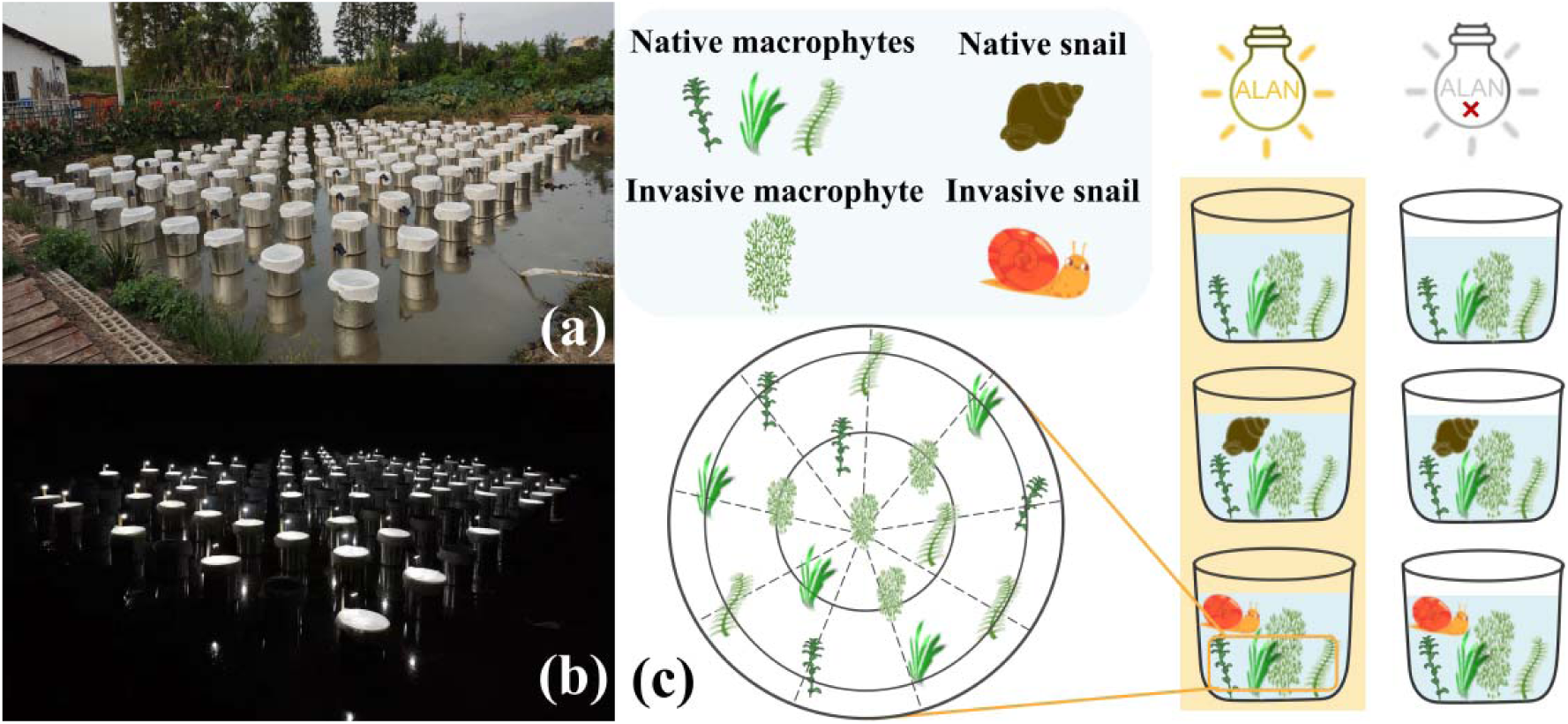
Schematic of Experiment 3, showing bucket mesocosms crossed with light (ALAN vs. No-ALAN) and herbivore (no snails, invasive *Pomacea canaliculata*, native *Cipangopaludina chinensis*) treatments. (**a**) Daytime photograph of the bucket arrangement with macrophytes. (**b**) Nighttime photograph illustrating the ALAN treatment with solar lamps. (**c**) Overview of the factorial design applied to macrophyte communities (*Vallisneria natans*, *Hydrilla verticillata*, *Myriophyllum spicatum*, *M. aquaticum*), with positions of individual macrophytes indicated. Photos in (**a**) and (**b**) were taken by Jingjing Xue. Illustrations of macrophytes, snails, and light bulbs in (**c**) were obtained from open-source material libraries in WPS Office and Microsoft PowerPoint and combined using PowerPoint.The experiment was perfomed at the Dongting Lake Wetland Ecosystem Observation and Research Station.

**Figure S4.**
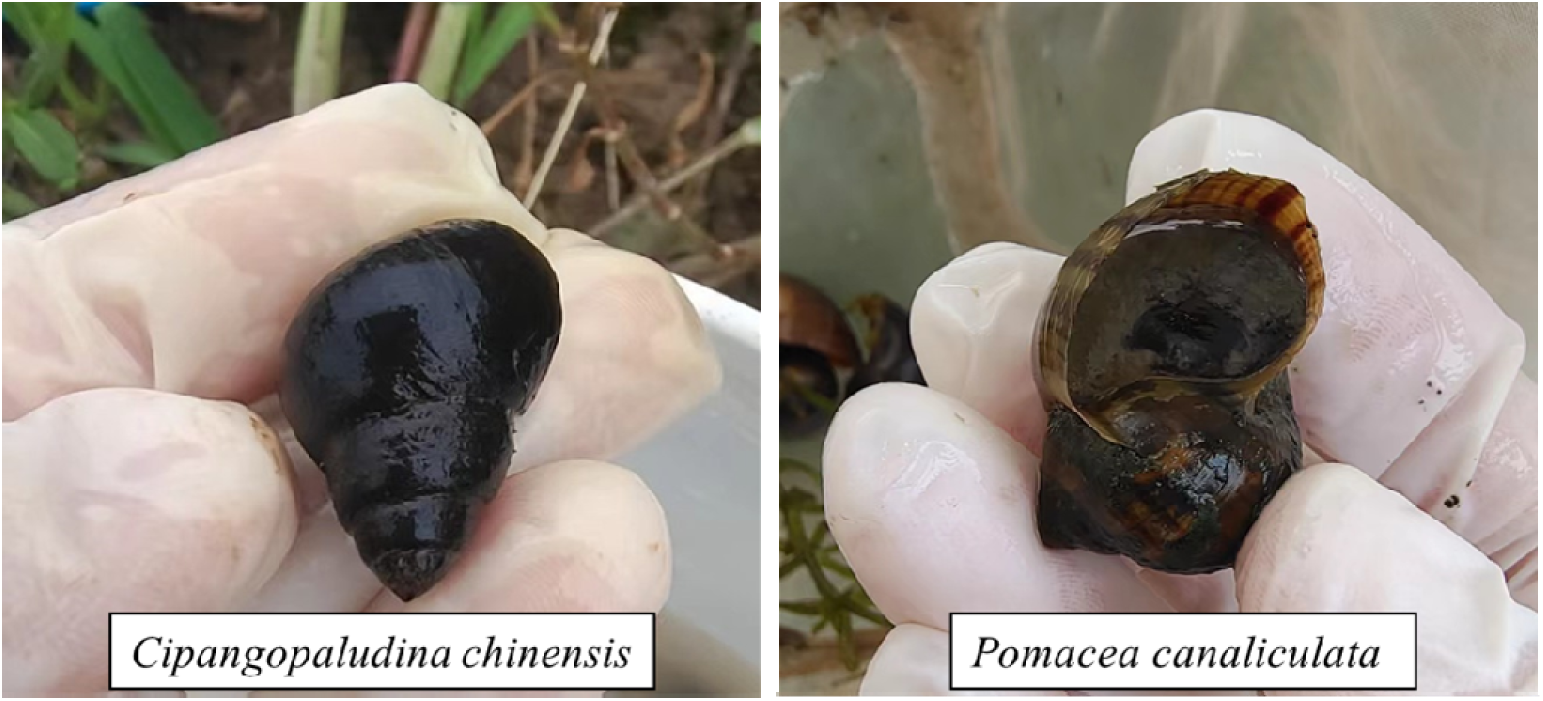
Specimens of the native snail *Cipangopaludina chinensis* (left) and the invasive snail *Pomacea canaliculata* (right), collected from ponds near the National Field Scientific Observation and Research Station of Dongting Lake Wetland Ecosystem, Hunan, China (29°30′N, 112°48′E). Photo credit: Jingjing Xue.

## Appendix S2

### Section S1: Methods

To test whether artificial light at night (ALAN) influences the incubation of *Pomacea canaliculata* eggs, we collected 40 egg masses attached to plant stems near the research station on August 19, 2023. Each stem was placed obliquely in a glass beaker containing 5 cm of groundwater, and the 40 beakers were randomly distributed across four outdoor cement ponds (1 m³) under a canopy, with half exposed to ∼30 lux ALAN from sunset to dawn. We recorded the number of hatched juveniles daily and ended the experiment on September 26 after three consecutive days with no new hatchlings. The total number of juveniles per beaker was calculated as the sum of daily counts. The native snail *Cipangopaludina chinensis* was not evaluated because its longer reproductive cycle (Stephen et al. 2013) precluded reproduction during the experiment.

To analyze treatment effects, we used a generalized linear mixed-effects model (*glmmTMB* package) (Magnusson et al., 2017) with an ‘nbinom1’ error distribution. Light treatment (ALAN vs. No-ALAN) was included as the fixed effect and total hatchlings per beaker as the response. To account for differences in initial egg numbers, we included the total egg count per mass (hatched + unhatched eggs) as an offset. Pond identity was included as a random effect, and significance of the fixed effect was assessed with log-likelihood ratio tests (Zuur et al. 2009).

### Section S1: Results

Hatching of invasive snail *Pomacea canaliculata* eggs began on August 22 and was initially unaffected by ALAN (Appendix S2: Fig. S1). Nevertheless, peak hatching occurred earlier under ALAN, with a maximum of 49.95 hatchlings emerging on August 29. Overall, ALAN significantly accelerated incubation, leading to a 28.6% increase in total hatchlings (Appendix S2: Fig. S1).

**Figure S1.**
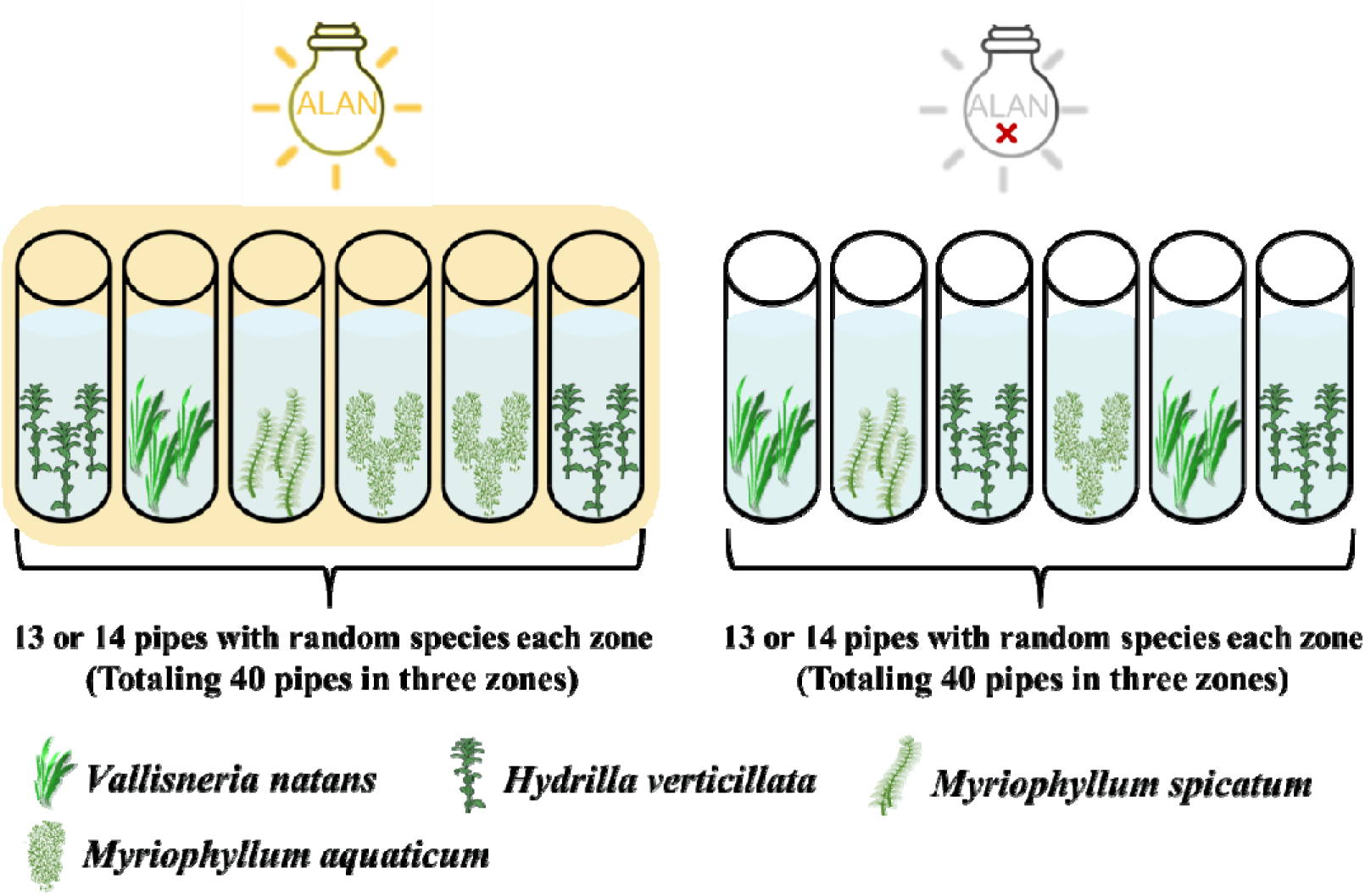
Daily mean (±1 SE) number of *Pomacea canaliculata* hatchlings under ALAN and No-ALAN treatments. The inset figure shows total hatchlings (±1 SE) under both light treatments (light treatment: d.f = 1, χ² = 5.225, *p* = 0.022). Each black dot represents one egg-mass sample.

